# Short-term BDD treatment of wastewater effluent reveals dissolved oxygen turnover and microbial community restructuring

**DOI:** 10.64898/2026.09.24.754035

**Authors:** Anna-Neva Visser, Manuel Zulla, Stefan Rosiwal, Andreas Burkovski

**Affiliations:** GeoZentrum Nordbayern, Friedrich-Alexander-Universität Erlangen-Nürnberg, Schlossgarten 5, 91054 Erlangen, Germany; Department of Material Sciences and Engineering, Friedrich-Alexander-Universität Erlangen-Nürnberg, Martensstr. 5, 91508 Erlangen, Germany; Department of Biology, Microbiology Division, Friedrich-Alexander-Universität Erlangen-Nürnberg, Staudtstr. 5, 91058 Erlangen, Germany

**Keywords:** Wastewater treatment plants, wastewater bacterial community, DIC/DOC/POC, DO, isotopes, shotgun metagenomic sequencing, EAOP, boron-doped diamond

## Abstract

Electrochemical advanced oxidation processes (EAOPs) are increasingly investigated for tertiary wastewater treatment, yet their short-term effects on real wastewater effluent remain poorly constrained. This study investigated real wastewater treatment plant (WWTP) effluent following short-term boron-doped diamond (BDD) electrochemical treatment using an interdigitated double-diamond electrode (iDDE), combined with hydrochemical, stable-isotope and shotgun metagenomic analyses. After 10 min at 7.0 V and 1.02 A, physicochemical parameters and bulk carbon pools remained largely unchanged. In contrast, δ^18^O-DO decreased by 31.3 ±0.28‰ alongside a small increase in dissolved oxygen (0.16 ±0.03 mmol/L), indicating substantial oxygen consistent with electrochemical oxygen generation. Shotgun metagenomics revealed pronounced restructuring of the recovered microbial DNA pool, including increases in *Comamonadaceae* (∼12.3% vs. ∼25.1%) and *Zoogloeaceae* (∼3.3% vs.∼8.4%), whereas the oxidative-stress-defence gene repertoire showed no coherent response. These taxonomic changes cannot be interpreted as selective survival as metagenomics do not resolve viability or intra-/extracellular DNA contribution. Overall, short-term iDDE treatment affected the microbial component of real WWTP effluent despite comparatively limited changes in bulk chemistry, highlighting that EAOP effects extend beyond conventional disinfection.

**Highlights:**

- An interdigitated double diamond electrode (iDDE) was tested on real WWTP effluent
- BDD treatment strongly decreased δ^18^O-DO despite minor changes in bulk chemistry
- Shotgun metagenomics revealed pronounced restructuring of the recovered microbial DNA
- Oxidative-stress-defence gene potential showed no coherent treatment response
- EAOP effects on wastewater microbiota extend beyond conventional disinfection

## 1. Introduction

Modern wastewater treatment plants (WWTPs) safeguard aquatic ecosystems and protect drinking-water resources through sequential physical, chemical and biological treatment processes (United Nations World Water Assessment Programme 2014; Pistocchi et al. 2019). Although conventional WWTPs efficiently reduce organic matter and nutrients (Bolisetty et al. 2019; Choi et al. 2024), they were not designed to eliminate the microbial fraction of wastewater. Consequently, treated effluent discharged into receiving water bodies not only affects existing microbial communities but also contributes to their microbial diversity (Kalinowska et al. 2022; de Santana et al. 2022; Dai et al. 2023). This contribution encompasses opportunistic pathogens and microorganisms carrying antimicrobial resistance genes (Varela and Manaia 2013; Liu et al. 2018; Osińska et al. 2020), making WWTPs recognised point sources for microbial and genetic contamination (Gao et al. 2022). To address contaminants escaping conventional treatment, increasing attention has been directed towards the implementation of an advanced “fourth treatment stage”(Eggen et al. 2014; Ianes et al. 2025; Di Cesare et al. 2026). Although this concept has primarily focused on chemical micropollutants (e.g., Svahn and Borg 2024; Busck et al. 2026), the microbial component of treated effluent represents an equally important target, particularly for energy-efficient and chemical-free treatment technologies (Norra et al. 2022; Kato and Kansha 2024; Schröder et al. 2026).

Among those novel treatment technologies, electrochemical advanced oxidation processes (EAOPs) have attracted increasing attention (Oliveira et al. 2025; Pasciucco et al. 2025; Zhong et al. 2025). At boron-doped diamond (BDD) electrodes, anodic water oxidation generates a mixture of reactive oxygen species (ROS), most notably hydroxyl radicals (.OH) (e.g., Kraft 2007). In contrast to active anodes, BDD favours the formation of weakly physisorbed .OH species rather than strongly chemisorbed oxygen species and higher oxidation states, thereby promoting highly reactive and comparatively non-selective oxidation (Kraft 2007; Sirés et al. 2014; Martínez-Huitle and Brillas 2021). These ROS can attack a broad range of cellular components including membrane lipids, proteins and nucleic acids, thereby rendering the harbouring cells inactive (Schorr et al. 2019; Hong et al. 2019; Seixas et al. 2022). However, conventional BDD systems are often energy intensive and exhibit limited performance in the low-conductivity matrices characteristic of treated municipal effluent (Ma et al. 2018). The recently developed interdigitated double diamond electrode (iDDE) represents a modified BDD electrode configuration designed to overcome limitations associated with conventional electrode geometries. Building on the earlier laser-structured double diamond electrode (DDE) design (Koch et al. 2020), the iDDE incorporates a substantially reduced interelectrode distance of <50 µm, thereby minimising ohmic losses and reducing energy demand by 14-46% during the oxidation of organic pollutants (Zulla et al. 2025). This closely spaced geometry further enabled direct BDD-based electrochemical treatment of comparatively low-conductivity matrices without requiring supporting electrolytes or proton exchange membranes (Zulla et al. 2025).

Yet, despite these technological advances, comparatively little is known about how electrochemical oxidation modifies the physicochemical environment of real wastewater effluent. While electrochemical reactions occurring at the electrode surface are well described (e.g., Einaga 2022), the chemical conditions established within the treated water are more difficult to characterize because the most reactive oxidants (i.e., ROS) are transient and challenging to quantify directly. Consequently, complementary approaches capable of tracing electrochemical processes within the treated matrix are required. Stable isotope analyses could offer such an opportunity. In particular, the stable isotope composition of water (δ^2^H/δ^18^O-H2O) and the dissolved oxygen (δ^18^O-DO), with the latter widely applied to trace oxygen sources and oxygen cycling in aquatic systems (e.g., Dordoni et al. 2022; Maier et al. 2025). Hence, δ^18^O-DO signatures therefore have the potential to reflect oxygen being generated through electrochemical water oxidation, providing an indirect tracer of anodic processes associated with BDD electrochemistry. In parallel, dissolved organic and inorganic carbon (DOC/DIC) and particulate organic carbon (POC), together with their respective stable carbon isotope composition (δ^13^C), could provide a broader assessment of whether short-term electrochemical treatment measurably alters bulk carbon pools and carbon speciation within the effluent. Combined with conventional physicochemical measurements, these complementary geochemical signatures may therefore provide valuable insight into the chemical environment generated during electrochemical treatment without relying on direct quantification of individual reactive species.

Similarly, the biological consequences of electrochemical oxidation within complex communities are not fully understood (Oluoch et al. 2024). Most studies evaluating electrochemical disinfection rely on culture-based measurements or a limited number of organisms (Valero et al. 2017; Anfruns-Estrada et al. 2017; Huo et al. 2022; Kumar and Kanmani 2022), providing estimates of overall microbial inactivation offering little information on changes occurring across the broader community. Amplicon sequencing has expanded community-level investigations but remains constrained by primer bias, variable rRNA gene copy numbers and limited functional resolution (Hackmann 2025). In contrast, shotgun metagenomic sequencing simultaneously characterises the taxonomic composition and functional gene repertoire of recovered community DNA, including genes associated with oxidative-stress defence and other cellular functions potentially affected by electrochemical oxidation. Although this approach does not distinguish DNA originating from viable and non-viable cells, it enables a community-level assessment of how the recovered microbial DNA pool changes following treatment.

Against this background the present pilot study explores whether integrating physicochemical analyses, stable isotope geochemistry and shotgun metagenomics can provide complementary insight into the consequences of short-term BDD-based electrochemical treatment of real wastewater effluent using an interdigitated double diamond electrode (iDDE). Specifically, this study aimed to (i) determine changes in the physicochemical, redox and carbon characteristics of treated effluent; (ii) evaluate whether dissolved oxygen stable isotope signatures (δ^18^O-DO) provide evidence consistent with electrochemical water oxidation, while assessing changes in bulk carbon pools and their isotope composition; and (iii) identify changes in the taxonomic and functional composition of recovered microbial DNA pool following BDD-based electrochemical treatment. By integrating these independent lines of evidence, this exploratory study provides an initial mechanistic framework for evaluating electrochemical wastewater treatment beyond conventional reactor performance metrics.

## 2. Material & Methods

### 2.1. Area of study and effluent provenance

On-site electrochemical testing was conducted at the wastewater treatment plant No. 1 (Klärwerk 1) in Nürnberg-Muggenhof, Germany (49.4620° N; 11.0470° E.). This mechanical-biological-chemical facility has a design capacity of 1,400,000 population equivalents (PE) and processes a mixed municipal and industrial influent via a two-stage activated sludge process configured for full nitrification, denitrification, and enhanced chemical phosphorus removal (Stadtentwässerung und Umweltanalytik Nürnberg (SUN) 2020; Ingenieurbüro Miller 2021). The biological treatment sequence is followed by a deep-bed wastewater filter (sand filtration) as the final cleaning step prior to discharge into the Pegnitz River (Stadtentwässerung und Umweltanalytik Nürnberg (SUN) 2020).

All measurements were performed within the facility’s Centralized Online Analytics and Control Centre (Zentrale Onlineanalytik), an integrated laboratory and monitoring complex. Here, continuous-flow split streams from every individual treatment stage are permanently pumped into a central analytical hall equipped with dedicated HACH sensor stations for real-time water quality monitoring. Fully treated final effluent was accessed and abstracted directly from the corresponding post-filtration sampling line within this hall, establishing the final matrix baseline for downstream evaluation by the interdigitated double diamond electrode (iDDE).

### 2.2. Interdigitated double diamond electrode (iDDE)

*iDDE* – Electrochemical exposure was performed using an iDDE, a BDD-based electrode configuration derived from the earlier laser-structured double diamond electrode design (Koch et al. 2020) and further developed for electrochemical treatment of low-conductivity waters (Zulla et al. 2025)(2025)(Koch et al. 2020). The iDDE features a planar boron-doped diamond (BDD) layer on a non-conductive ceramic substrate, structurally modified via laser ablation to achieve an interelectrode spacing (IES) <50 µm. This microscale BDD geometry facilitates highly energy-efficient electrochemical advanced oxidation processes (EAOPs) in low-conductivity matrices, enabling direct treatment without the need for supporting electrolytes or proton exchange membranes (Zulla et al. 2025).

*Experimental setup and electrochemical treatment* – Effluent water was collected as sequential grab samples (*n* = 3) from the designated sampling station at the Centralized Online Analytics and Control Centre. Samples were either processed immediately to establish the baseline condition (untreated controls) or subjected to electrochemical advanced oxidation using the interdigitated double diamond electrode (iDDE; see Fig. 1). For the treated setup, 2 L of effluent water were transferred into a clean plastic beaker and placed on a magnetic stirring plate (600 rpm, IKA). The iDDE was immersed in the matrix and operated at a constant cell voltage of 7.0 V and a current of 1.02 A under continuous, uniform magnetic stirring for exactly 10 min. Immediately after the 10 min treatment period, both the stirring mechanism and the electrical power supply were terminated. In-field physicochemical parameters were determined *in situ*. Subsequently, distinct sample aliquots were taken and partitioned for downstream analyses as described below.

**Figure 1:**
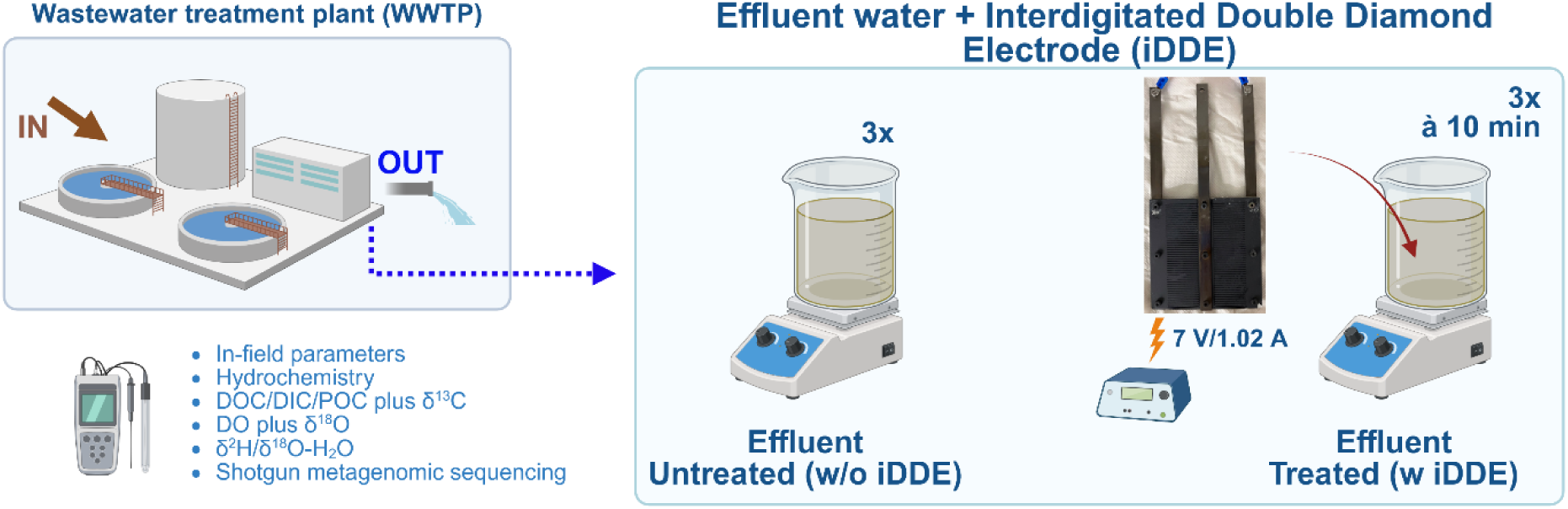
Experimental overview of the wastewater effluent treatment and analytical framework. Sequential grab samples (*n* = 3) were collected from the designated sampling station at the Centralized Online Analytics and Control Centre in the WWTP. The electrochemical advanced oxidation setup consisted of untreated controls and treated effluent water exposed to an interdigitated double diamond electrode (iDDE) for 10 min at a constant voltage of 7.0 V and current of 1.02 A under continuous magnetic stirring. In-field parameters were determined *in situ* and samples were taken to determine potential changes in hydrochemistry, carbon pools (dissolved organic/inorganic carbon, particulate organic carbon and their respective δ^13^C values), dissolved oxygen and δ^18^O-DO, as well as water isotopes (^2^H/δ^18^O-H2O), and shotgun metagenomic profiles.

### 2.3. Water chemistry

*Sample collection* – Similar to water collected for molecular analyses, effluent was collected in 2 L plastic beakers. Subsamples were taken to measure water pH, temperature (°C), electric conductivity (EC), redox potential (Eh), and oxygen (O2) using on-site calibrated multiparameter sensors (Multi 3620 IDS, WTW GmbH). Total alkalinity was determined on-site via acid-base titration using a hand-held, digital titrator and 1.6 N H2SO4 cartridges (HACH). For laboratory analyses, effluent was immediately filtered (0.45 µm PES, Sartorius AG, Minisart High-Flow) in triplicate into the respective vials. Samples for major ion and trace element analyses were collected in 15 mL centrifugation tubes (BD Biosciences). Samples for δ^18^O/δ^2^H-H2O were collected in 12 mL exetainer vials (LABCO). Samples for δ^18^O-DO analyses were collected in identical exetainers, but pre-poisoned with 20 µl saturated HgCl2. Samples for dissolved inorganic/organic carbon (DIC/DOC), and their corresponding δ^13^C isotope values, were collected in 40 mL amber glass vials pre-poisoned with 20 µl saturated HgCl2. For POC determination, samples were vacuum-filtered (see 2.4) until the filters clogged, transferred into petri dishes using clean stainless-steel tweezers, and stored at 4°C until further processed. All vials were sealed with Parafilm and stored at 4 °C in the dark until analysis.

*Major ion and trace metal quantification* – Effluent samples filtered for major cation and trace element analysis were acidified to pH <2 with 65% HNO3 (suprapur) and quantified using an ICP-MS (iCAP Qc, ThermoFisher; precision better than 1 % (1σ)). Unacidified aliquots were analysed for major anions by ion chromatography (Compact IC Flex, Metrohm; precision better than 5 % (1σ)). Details are provided in Visser et al. (2026).

### 2.4. Isotopes

*Carbon species quantification and isotope analyses* – Dissolved inorganic and organic carbon (DIC and DOC) concentrations, as well as their corresponding isotope ratios (δ^13^C-DIC and δ^13^C-DOC), were determined using an OI Analytical Aurora 1030W TIC/TOC analyser coupled in continuous flow mode to a Thermo Fisher Scientific Delta V Plus isotope ratio mass spectrometer (IRMS). By reacting the sample with 1 mL of 5% H3PO4 at 70 °C for 2 min, the DIC was liberated as CO2. DOC was oxidised to CO2 on a separate sample aliquot via reaction with 2 mL of 10% Na2S2O8 at 98 °C for 5 min. For both, the produced CO2 was purged from solution by helium, passed through a Nafion trap and quantified using non-dispersive infrared (NDIR), and transferred to the IRMS for isotope analyses. For low concentrations, a trap and purge (T&P) system was installed. Details addressing the coupling between TIC/TOC analyser and IRMS are published in St-Jean (2003).

For POC determination, effluent water was filtered using pre-weighted glass fibre filters (GF-5, 0.4 µm; Macherey-Nagel), that were pre-combusted at 400°C for 4 h to remove any residual carbon. For analysis, filters were treated as described by Maier et al. (2025) and both carbon contents and the ^13^C/^12^C isotope ratios were measured using an elemental analyser (Costech ECS 4010, NC Technologies) coupled in helium continuous-flow mode to an IRMS (Delta V, ThermoFisher).

All δ^13^C values are expressed in the standard δ-notation (‰) relative to Vienna Pee Dee Belemnite (VPDB):

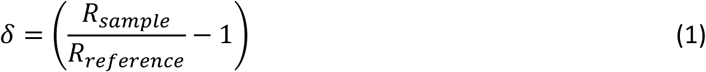

where *R* is the *n*(^13^C)/*n*(^12^C) ratio in the sample and reference (Coplen 2011). Data were corrected for instrumental drift and linearity following van Geldern et al (2013). Repeat analyses of standards (KHP, C3 and C4 sugars) yielded precisions better than ±0.3‰ (1σ).

*Dissolved oxygen and water isotopes* – δ^18^O of dissolved oxygen (DO) was determined using a modified automated equilibration system (Gasbench II, ThermoFisher) coupled in continuous-flow mode to an IRMS (Delta V Advantage, ThermoFisher), following the procedures of Barth et al. (2004) and Wassenaar and Koehler (1999), with further methodological details given in Maier et al. (2025). Briefly, DO extraction was conducted by equilibrating each water sample with a 3 mL helium headspace through vigorous shaking; the equilibrated headspace was then analysed for both DO concentration and δ^18^O. All samples were analysed in triplicate, with external reproducibility better than ±0.2 ‰ (1σ). δ^18^O and δ^2^H of water (δ^18^O-H2O, δ^2^H-H2O) were measured using cavity ring-down spectroscopy (CRDS) on a Picarro laser analyser (L1102-*i* WSCRDS), following the protocol of van Geldern and Barth (2012). Values are reported in the δ-notation as defined above (Eq. 1), relative to the Vienna Standard Mean Ocean Water (VSMOW; Baertschi 1976). External reproducibility was better than ±0.1 ‰ and 1% (1σ) for δ^18^O and δ^2^H, respectively.

### 2.5. Molecular biology-based methods

*Biomass collection* – Effluent water was collected at the designated station at the Centralized Online Analytics and Control Centre using a 2 L plastic beaker. Sequential grab samples (*n* = 3) were collected and immediately vacuum-filtered through polyethersulfone membrane filters (0.2 µm, PES; Sartorius) using an autoclaved filtration tower (PS, Nalgene). Following filtration, using sterilized tweezers, filters were immediately transferred into sterile, RNase-/DNase-free 15 mL centrifugation tubes (VWR), placed in a temporary cooling box maintained at ∼-10°C for a maximum of 8 h during field transit, and subsequently stored at-80°C until downstream processing. For iDDE-treated samples, the effluent water was exposed to the iDDE as detailed in Section 2.2 prior to undergoing the identical filtration and storage protocol described above.

*DNA extraction and quality control* – Microbial genomic DNA was extracted from the frozen filters using the DNeasy PowerBiofilm Kit (Qiagen) according to the manufacturer’s instructions, with specific modifications to the mechanical lysis step to ensure complete biomass recovery from the resilient PES filter matrices. Therefore, filters were transferred directly into bead tubes and subjected to rather aggressive mechanical disruption via a FastPrep-24 homogenizer (MP Biomedicals), using 3 to 4 cycles of 30 s at a vertical velocity of 7.0 m/s. To mitigate frictional heating and prevent thermal degradation of the raw nucleic acids, samples were rapidly chilled on ice between consecutive bead-beating cycles. Total DNA was eluded in 100 µL of elution buffer, and its structural integrity was assessed via 1% (w/v) agarose gel electrophoresis in 1x TAE buffer. Therefore, samples were prepared using Purple Gel Loading Dye (6x, New England Biolabs), and fragments were sized relative to a peqGOLD DNA Ladder Mix (VWR). The gel was stained with ethidium bromide and visualized under UV transillumination. To confirm that the extracts met the strict quantification and purity requirements for subsequent shotgun metagenomic sequencing, fluorometric quantification was performed using the Qubit dsDNA BR Assay Kit (Invitrogen; see SI Table 1).

*Shotgun metagenomic sequencing and primary bioinformatics* – The final DNA extracts were sent to CeGaT (Tübingen, Germany), who performed library preparation, sequencing, and primary bioinformatic assignment. Briefly, metagenomic libraries were prepared using the Illumina DNA Prep with (M) Tagmentation kit (Illumina) with an initial DNA input of up to 100 ng per sample (or lower for low-yield extracts). Sequencing was executed on the Illumina NovaSeq X Plus platform in paired-end mode with a targeted read length of 2×100 bp, generating ∼2 Gb (10×10^6^ clusters) per sample. Across all samples, 93.3 % of the sequenced bases achieved a Phred quality score ≥ 30 (Q30).

Base quality was monitored using FastQC on the Illumina DRAGEN Bio-IT Platform (v4.4.4), and a comprehensive quality report was compiled using MultiQC (v1.22.2). Raw sequence reads were demultiplexed with *bcl2fastq* (v2.20; Illumina) and adapter-trimmed via *Skewer* (v0.2.2; Jiang et al. 2014); no additional quality trimming was performed. General statistics provided by CeGaT are in SI Table 2. For taxonomic and functional profiling, a subset of 10 million adapter-trimmed forward reads (R1) per sample was aligned against the filtered NCBI RefSeq protein database (release 94) using DIAMOND in BLASTX mode (Buchfink et al. 2015). Aligned reads were taxonomically assigned utilizing the lowest common ancestor (LCA) algorithm in MEGAN6 Ultimate Edition (v6.18.7; Huson et al. 2016), filtering for taxa with relative sequence abundance >0.01 %. Functional annotation was simultaneously performed within MEGAN6 by mapping reads to KEGG orthology identifiers, alongside SEED, VFDB, and InterPro databases. Per-sample read counts and relative abundances stratified by taxonomic rank and functional hierarchy (KEGG levels 1-4) were provided for downstream analysis (see Section 3.4).

### 2.6. Data Analyses

Given the pilot scale of this study (two conditions, three sequential grab samples from the WWTP central collection point), the samples capture short-term temporal and technical variability rather than independent biological replication. As the available replication does not support robust statistical inference, inferential statistical testing was not applied. Hydrochemical and isotope data are presented as mean ±SD, and metagenomic community and functional data differences are reported descriptively. As these are relative sequence abundances derived from a compositional dataset, they describe shifts in community composition and metagenomic potential rather than changes in absolute abundance or gene expression.

Data clean up and visualisation were performed using SigmaPlot (v.9) and R (RStudio v2026.07.1+147; R Core Team 2014) using packages *ggplot2* (Wickham 2016), *dplyr* (Wickham et al. 2026); *tidyr* (Wickham et al. 2025b); *stringr* (Wickham 2025); and *scales* (Wickham et al. 2025a).

## 3. Results

### 3.1. BDD-induced changes in the physicochemical environment

Treatment with the iDDE (10 min, 7 V, 1.02 A) resulted in minor changes in the physicochemical parameters of the effluent (Fig. 2). Among the on-site measured parameters, redox potential (Eh) showed the most pronounced response, decreasing by 509.13 ±16.75 mV. In contrast, electrical conductivity (EC) exhibited a comparatively small decrease (ΔEC:-18.33 ±12.14 µS/cm), whereas pH, oxygen (O2) and temperature increased marginally (ΔpH: +0.13 ±0.02; ΔO2: +0.11 ±0.01 mmol/L; ΔT: +1.6 ±0.5°C). Alkalinity remained essentially unchanged (ΔAlk: +0.01 ±0.16 mmol/L).

**Figure 2:**
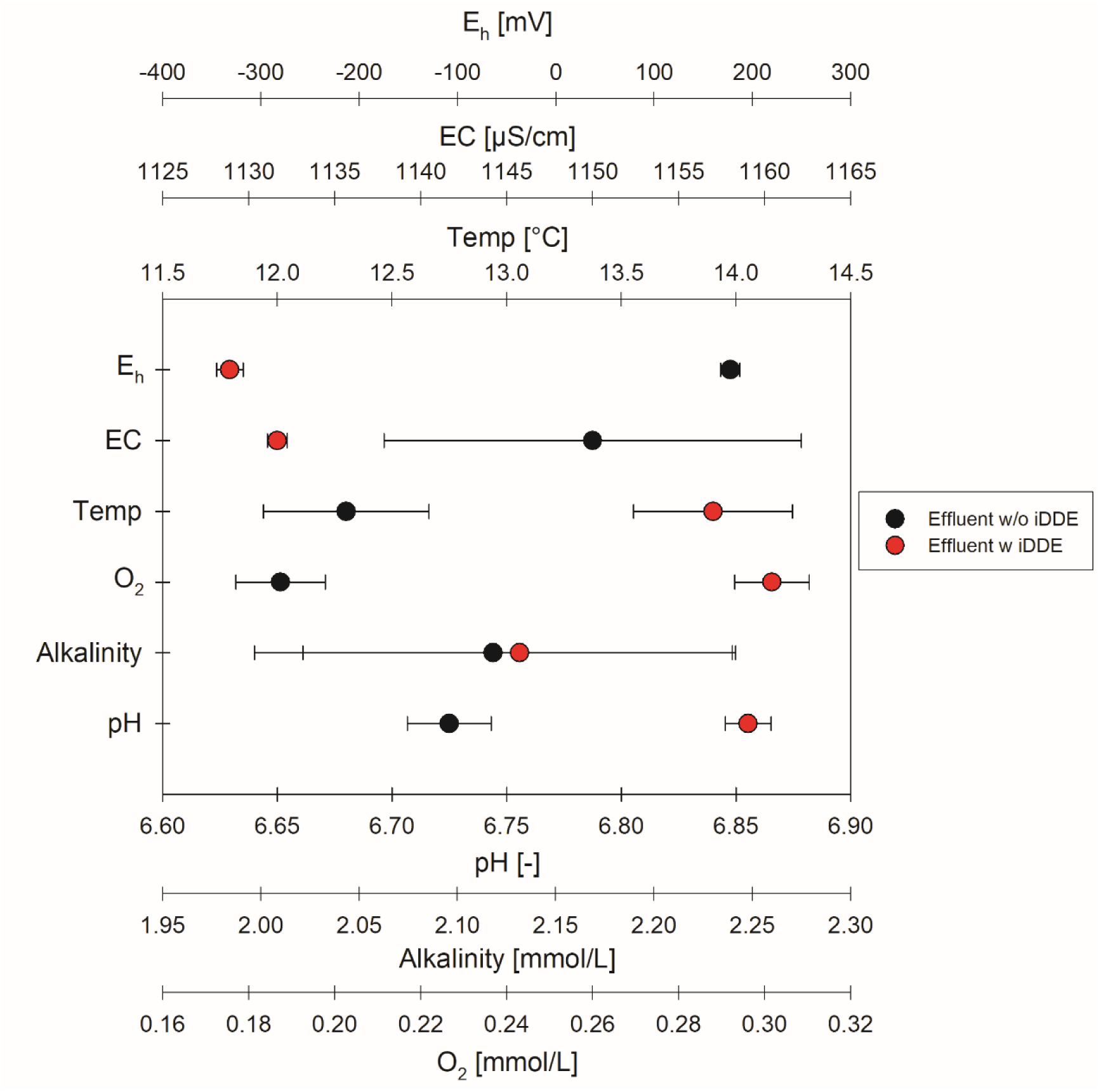
On-site physicochemical parameters (Multi 3620 IDS, WTW GmbH) and alkalinity (acid-base titration, 1.6 N H2SO4 cartridges (HACH)) for effluent treated with (red) and without (black) the interdigitated double diamond electrode (iDDE). Treated samples were exposed to the iDDE for 10 min (7 V, 1.02 A). Note different scaling of the x-axes. All values are also provided in ESM A. All values are means of *n* = 3 sequential grab samples. Error bars represent ±SD.

Concentrations of the major dissolved ions were largely unaffected by iDDE treatment (Fig. 3). Chloride, nitrate and sulphate concentrations remained unchanged (Fig. 3 A), while major cations Na, Mg, K, and Ca decreased only minor slightly (ΔNa: 0.08 ±0.11, ΔMg: 0.05 ±0.03, ΔK: 0.02 ±0.02, and ΔCa: 0.04 ±0.05 mmol/L; Fig. 3 B). Among the remaining analytes, phosphate decreased by 0.32 ±0.05 µmol/L (Fig. 3 A), while dissolved Mn and Fe exhibited the largest decreases, by 0.33 ±0.03 and 1.28 ±0.39 µmol/L, respectively (Fig. 3 B). Fe also showed the greatest variability among untreated samples (2.38 ±0.37 µmol/L). Concentrations of all additional measured elements are provided in ESM B.

**Figure 3:**
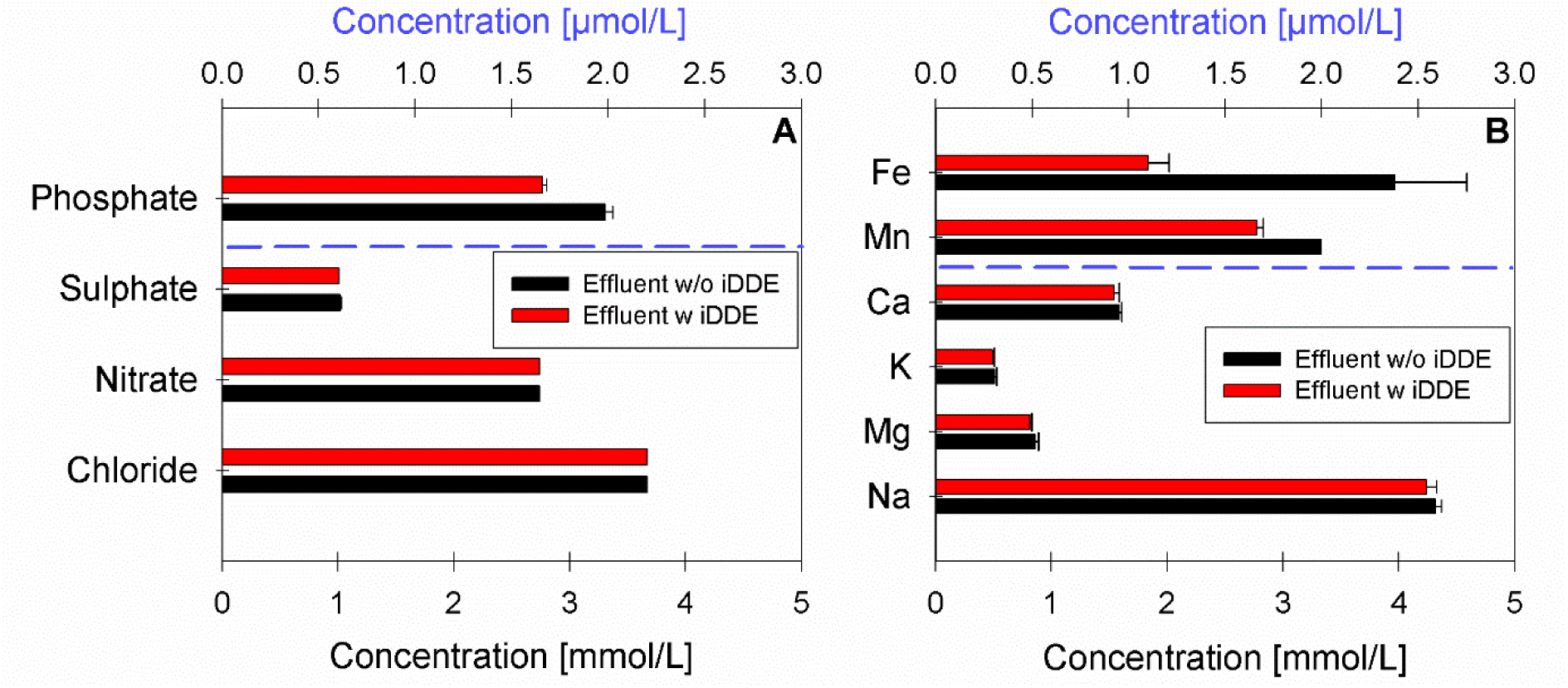
Concentrations of selected ions in effluent treated with (red) and without (black) the interdigitated double diamond electrode (iDDE). Treated samples were exposed to the iDDE for 10 min (7 V, 1.02 A). (A) Phosphate and the major anions (chloride, nitrate, sulphate); (B) major cations (Na, Mg, K, Ca) and the redox-active metals Mn and Fe. In both panels, species above the blue dashed line are given in µmol/L (upper axis) and species below the line in mmol/L (lower axis). Concentrations of all further measured elements are provided in ESM B. All values are means of *n* = 3 sequential grab samples. Error bars represent ±SD.

### 3.2. Carbon pool response to BDD-based electrochemical treatment

The carbon pool likewise exhibited only minor response to iDDE treatment (Fig. 4). Dissolved inorganic carbon (DIC) decreased by 0.10 ±0.14 mmol/L (Fig. 4 A), whereas dissolved organic carbon (DOC) changed only marginally (ΔDOC:-0.01 ±0.03 mmol/L; Fig. 4 B). Particulate organic carbon (POC) increased slightly (ΔPOC: +0.08 ±0.11 mg/L; Fig. 4 C). Stable carbon isotope compositions showed similarly small changes, with δ^13^C-DIC and δ^13^C-DOC decreasing by-0.16 ±0.11‰ and-0.08 ±0.03‰, respectively, while δ^13^C-POC increased by +0.21 ±0.19‰.

**Figure 4:**
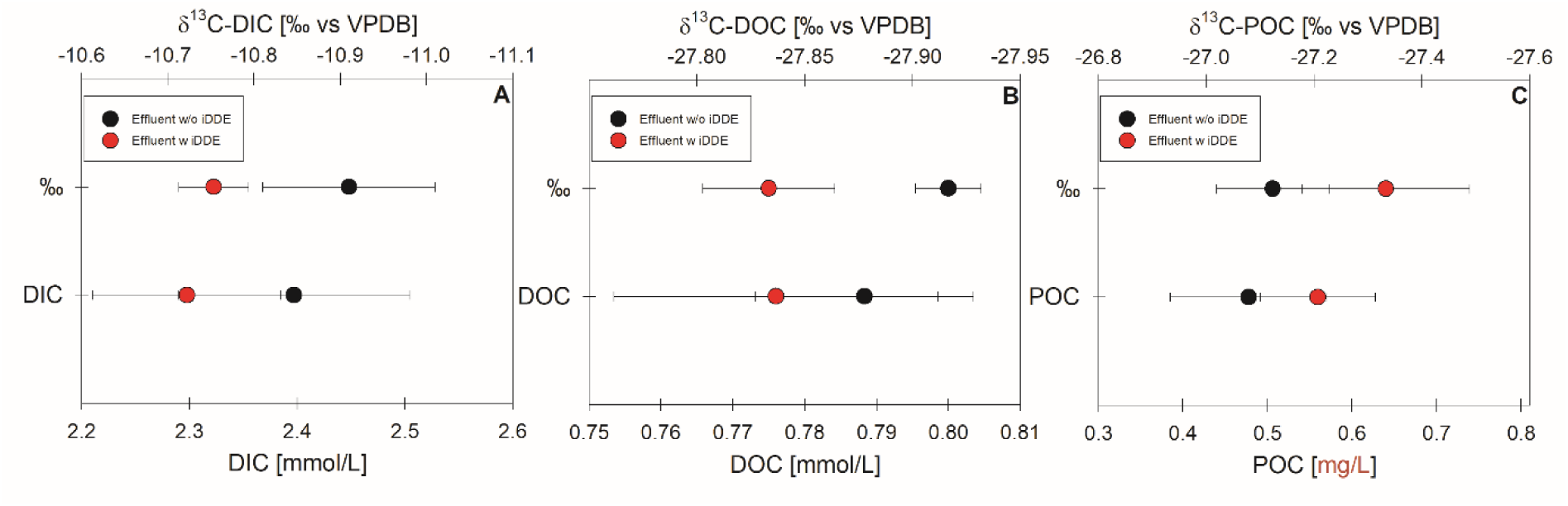
Concentrations and stable carbon isotope (δ^13^C) values of (A) dissolved inorganic carbon (DIC), (B) dissolved organic carbon (DOC), and (C) particulate organic carbon (POC) measured in effluent treated with (red) and without (black) the interdigitated double diamond electrode (iDDE). Treated samples were exposed to the iDDE for 10 min (7 V, 1.02 A). Note different scaling of the x-axes. All values are means of *n* = 3 sequential grab samples. Error bars represent ±SD. All values are also provided in ESM C.

### 3.3. Dissolved oxygen response to BDD-based electrochemical treatment

The isotopic composition of the DO pool was substantially altered in effluent treated with the iDDE (Fig. 5). Although DO concentrations calculated from δ^18^O-DO were slightly lower than the corresponding sensor measurements (w iDDE:-0.04 ±0.03; w/o iDDE:-0.08 ±0.02 mmol/L), both approaches showed a general increase in dissolved oxygen following treatment (ΔDO: +0.16 ±0.03 mmol/L), accompanied by a decrease in δ^18^O-DO of-31.3 ±0.28 ‰ (Fig. 5 A, B). In contrast, δ^2^H-H2O (Δ:-0.1 ±0.3 ‰) and δ^18^O-H2O (Δ: +0.05 ±0.07‰) were basically indistinguishable between treated and untreated effluent (Fig. 5 C).

**Figure 5:**
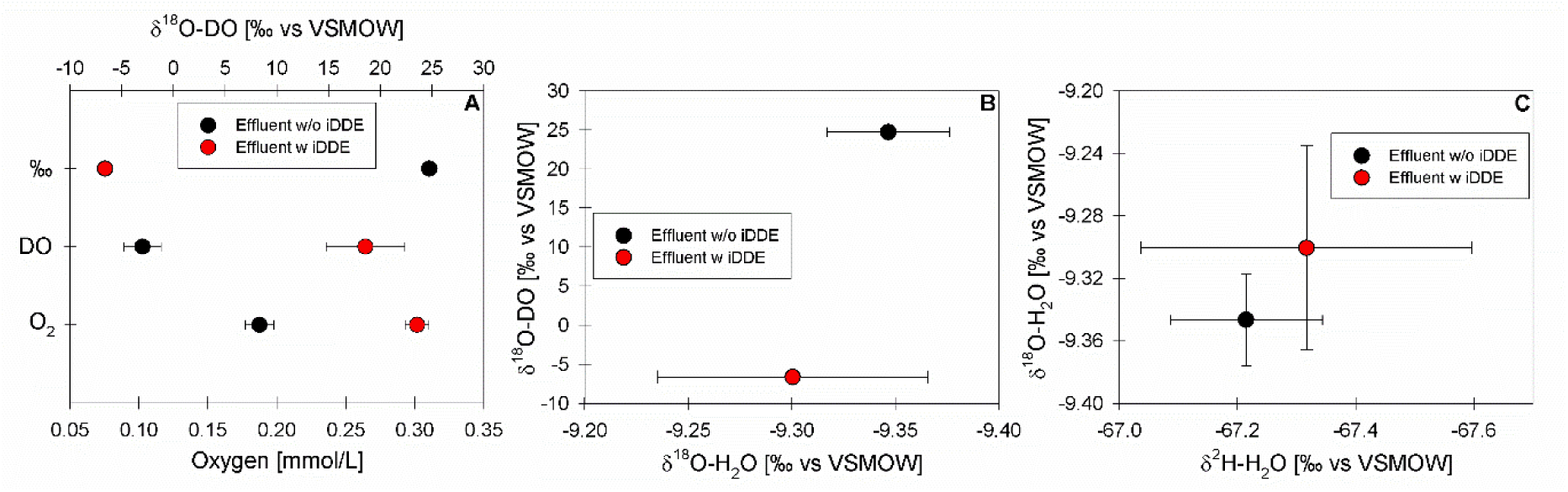
Dissolved oxygen and water isotope composition of effluent treated with (red) and without (black) the interdigitated double diamond electrode (iDDE). (A) Dissolved oxygen (DO) concentrations determined by optical sensor (O2) and calculated from δ^18^O-DO measurements (DO), together with corresponding δ^18^O-DO values; (B) Dual-isotope composition of the dissolved oxygen (δ^18^O-DO vs δ^18^O-H2O). (C) Stable isotope composition of water (δ^18^O-H2O vs δ^2^H-H2O). Note different scaling of the y-and x-axes. All values are means of *n* = 3 sequential grab samples. Error bars represent ±SD. All values are provided in ESM D.

### 3.4. BDD-based electrochemical treatment restructures the recovered microbial DNA pool

Shotgun metagenomic sequencing of six effluent grab samples yielded reproducible family-level taxonomic profiles, with iDDE-treated (WWTP_4,8,12) and untreated (WWTP_2,6,10) replicates clustering by treatment (Fig. 6). Between 59.5% (untreated) and 67.2% (treated) of sequence reads were assigned to the family level, while the remaining reads were left unclassified and were retained in all figures.

**Figure 6:**
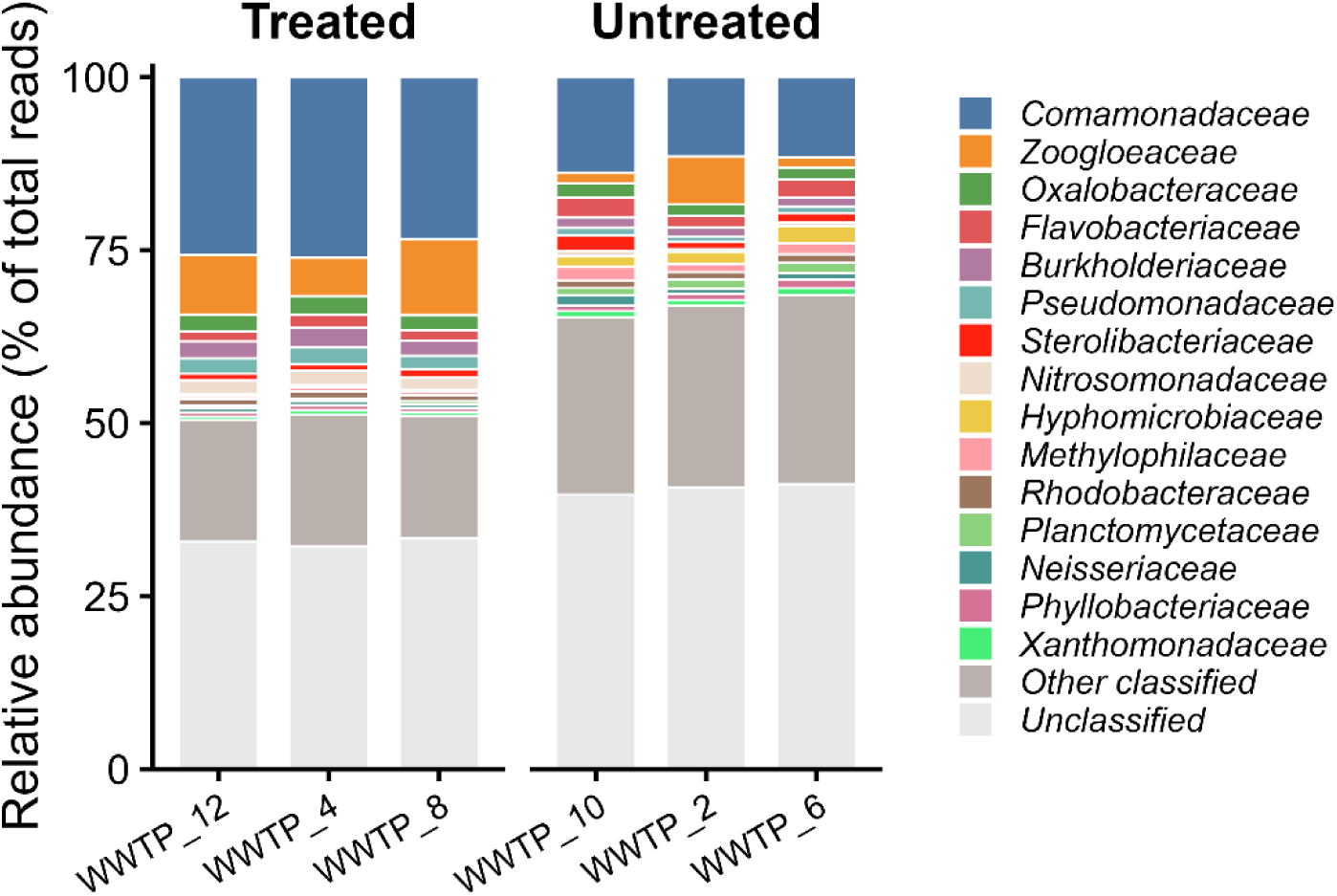
Relative sequence abundance of bacterial families in WWTP effluent with (treated; WWT_4,8,12) and without (untreated; WWTP_2,6,10) a 10 min interdigitated double diamond electrode (iDDE) treatment. Each bar represents one sequential grab sample. The 15 most abundant families across all samples are shown individually; all remaining classified families are grouped as “Other classified”, and reads that could not be assigned at the family level are shown as “Unclassified”. Relative abundances are expressed as percentage of total reads. The unclassified fraction is retained rather than normalized, so bars sum to 100%. Families are ordered by overall mean abundance.

Overall, iDDE-treatment was associated with shifts in the relative abundance of taxonomically assigned sequence reads (Fig. 6). The largest change was observed for *Comamonadaceae*, whose relative abundance increased from 12.3% in untreated samples to 25.1% following treatment. *Zoogloeaceae* likewise increased from 3.3% to 8.4%, although untreated replicates exhibited greater variability. Smaller increases were observed for *Nitrosomonadaceae*, *Pseudomonadaceae*, and *Burkholderiaceae*. Conversely, the relative abundance of *Hyphomicrobiaceae* (1.9% to 0.3%), *Methylophilaceae* (1.6% to 0.5%), *Planctomycetaceae* (1.3% to 0.4%), and *Microbacteriaceae* (1.2% to 0.3%) decreased following treatment.

At the genus level, 45.1% (untreated) to 54.5% (treated) of sequence reads were assigned, while the remaining reads were left unclassified (Fig. 7). Consistent with the family-level profiles, iDDE treatment was associated with reproducible shifts in the relative abundance of taxonomically assigned sequence reads. Within the *Comamonadaceae*, relative abundances increased for several genera, including *Acidovorax* (2.4% to 5.6%), *Malikia* (0.9% to 3.7%), *Variovorax* (0.6% to 2.2%), and *Rhodoferax* (0.9% to 1.9%). The largest increase at the genus level was observed for *Zoogloea* (2.2% to 5.6%; *Zoogloaceae*), while *Pseudomonas* (*Pseudomonadaceae*) increased from 0.9% to 2.2%. In contrast, the relative abundances of *Hyphomicrobium* (*Hyphomicrobiaceae*; 1.4% to 0.1%), *Methylotenera* (*Methylophilaceae*; 0.9% to 0.3%), *Flavobacterium* (1.4% to 0.8%), *Undibacterium* (0.9% to 0.5%), and *Hydrogenophaga* (*Comamonadaceae*; 0.8% to 0.4%) were lower following treatment.

**Figure 7:**
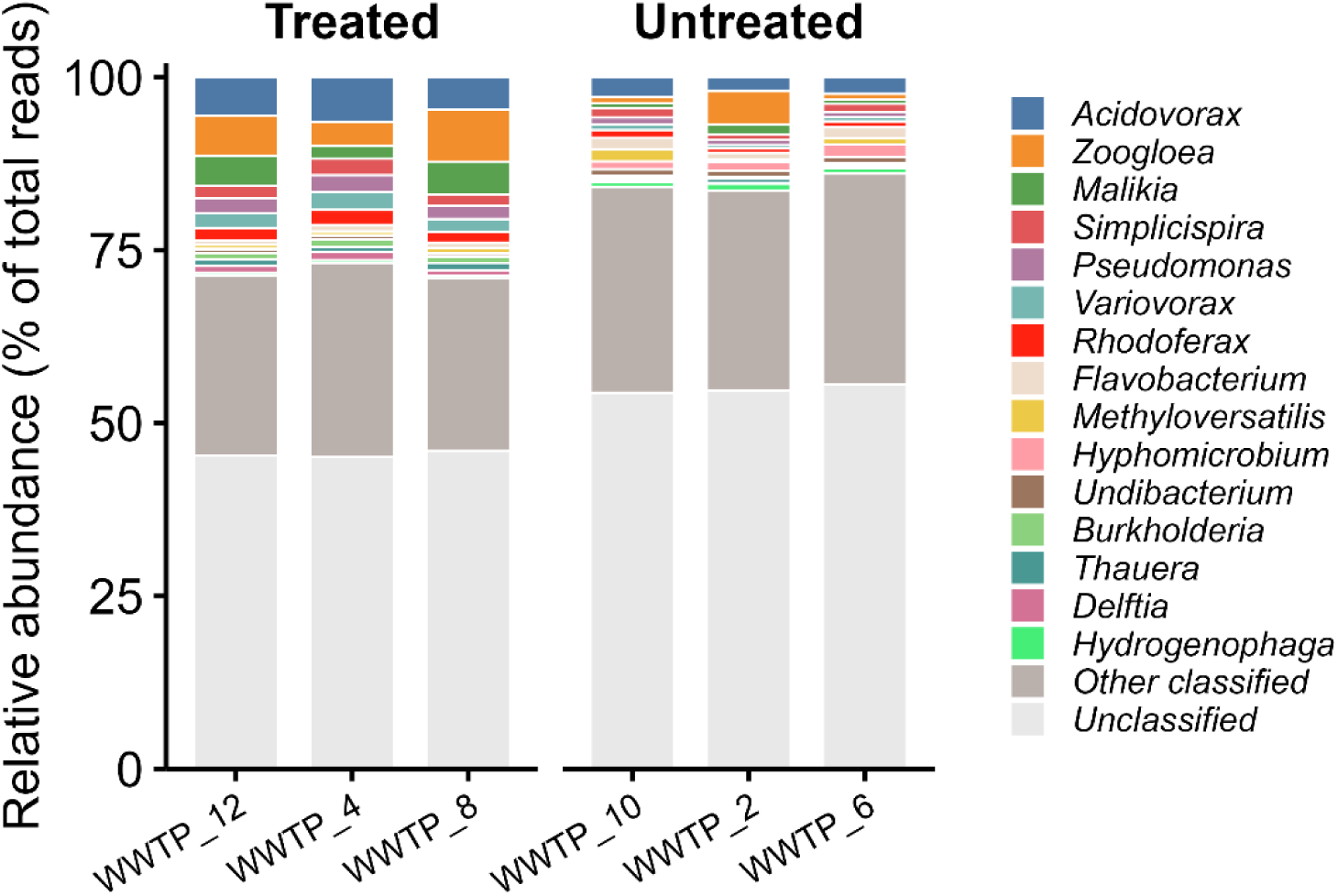
Relative sequence abundance of bacterial genera in WWTP effluent with (treated; WWT_4,8,12) and without (untreated; WWTP_2,6,10) a 10 min interdigitated double diamond electrode (iDDE) treatment. Each bar represents one sequential grab sample. The 15 most abundant genera across all samples are shown individually; all remaining classified genera are grouped as “Other classified”, and reads that could not be assigned at the genus level are shown as “Unclassified”. Relative abundances are expressed as percentage of total reads. The unclassified fraction is retained rather than normalized, so bars sum to 100%. Genera are ordered by overall mean abundance.

To identify the specific taxa driving these compositional shifts, eight families (Fig. 8 A) and eight genera (Fig. 8 B) exhibiting the greatest changes in mean relative abundances following iDDE-treatment were analysed. The direction of response was generally consistent across all three treated (w iDDE) grab samples relative to the untreated (w/o iDDE) controls. However, the response magnitude varied, and certain taxa (e.g., *Zoogloeaceae*) exhibited greater variability among the untreated samples. At the family level, relative abundance increases were most pronounced for *Comamonadaceae* and *Zoogloeaceae*. Conversely, at the genus level, the enrichment of *Comamonadaceae* was distributed across multiple constituent genera (i.e., *Acidovorax*, *Malikia*, *Variovorax*, *Rhodoferax*, and *Simplicispira*) rather than being dominated by a single taxon. Notable decreases in relative abundance occurred within families *Hyphomicrobioaceae* (*Hyphomicrobium*), *Methylophilaceae* (*Methylotenera*), and *Flavobacteriaceae* (*Flavobacterium*). These data reflect changes within the recovered DNA pool and do not necessarily imply shifts in absolute abundance or cell viability.

**Figure 8:**
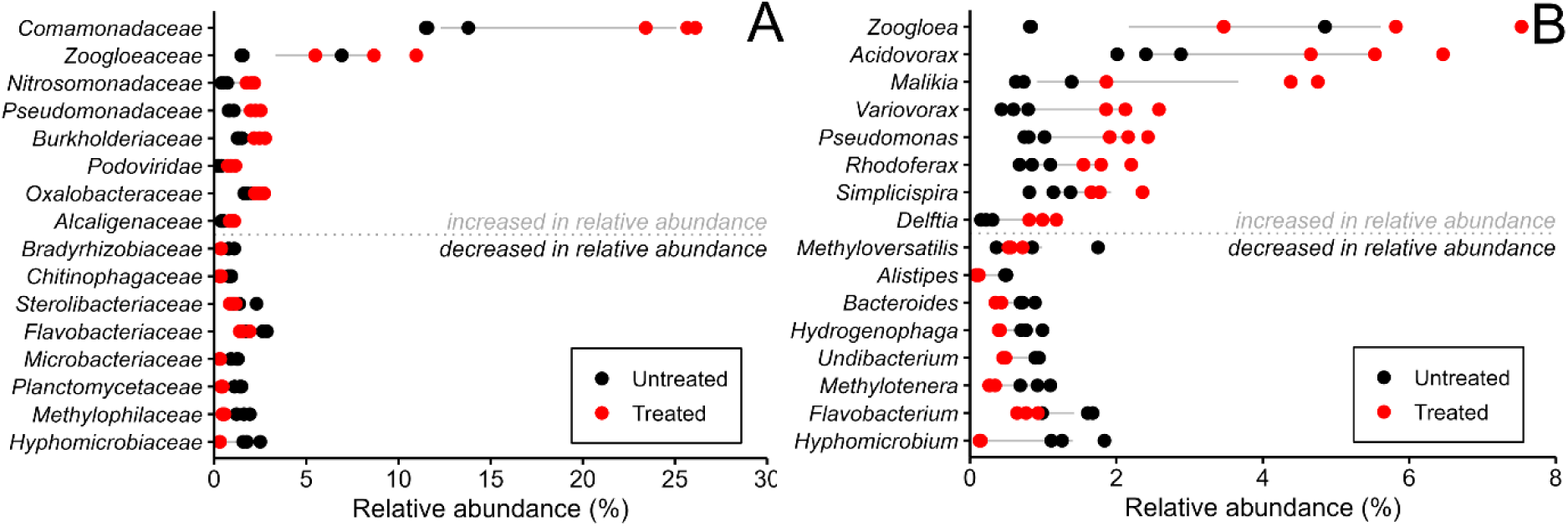
Bacterial taxa showing the largest differences in shotgun metagenomic DNA-derived relative sequence abundance between untreated (black) and iDDE-treated (red) wastewater treatment plant effluent, at the family (A) and genus (B) level. For each panel, the eight taxa with the largest increase and the eight with the largest decrease in mean relative abundance following treatment (treated – untreated) are shown, ordered by the magnitude and direction of that difference; the dotted grey line separates taxa that increased (above) from those that decreased (below) in relative abundance. Each point represents one sequential grab sample (*n* = 3 per condition), and the grey line connects the group means. Note that the x-axis ranges differ between panels to accommodate the higher family-level abundances. Relative sequence abundances describe the relative representation of taxa within the recovered DNA pool; they do not reflect absolute abundance or cell viability.

The relative abundances of 18 KEGG orthologs (KEGG level 4) encoded within the community metagenome and associated with oxidative stress defence were compared between the untreated and iDDE-treated samples to assess shifts in the metagenomic potential (Fig. 9). Differential abundance is expressed as the change in mean relative abundance (treated – untreated) in percentage points. Among the 18 profiled KEGG orthologs, six were higher and 12 were lower in treated samples. Yet, all shifts fell within a narrow range of approximately-0.055 to +0.005 percentage points. The enriched functional features included thioredoxin 2, alkyl hydroperoxide reductase subunit F (AhpF), cytochrome *c* peroxidase, the hydrogen peroxide regulator OxyR, glutathione peroxidase, and thiol peroxidase. However, their individual increases were marginal, remaining below 0.005 percentage points. Conversely, depleted KEGG orthologs included thioredoxin reductase, catalase-peroxidase (KatG), Fe-Mn superoxide dismutase, thioredoxin 1, peroxiredoxin Q, and catalase. The most substantial depletion was observed for thioredoxin reductase, which decreased by approximately - 0.055 percentage points. Importantly, these differences reflect changes in genomic capacity within the recovered DNA pool rather than active transcriptional or metabolic responses.

**Figure 9:**
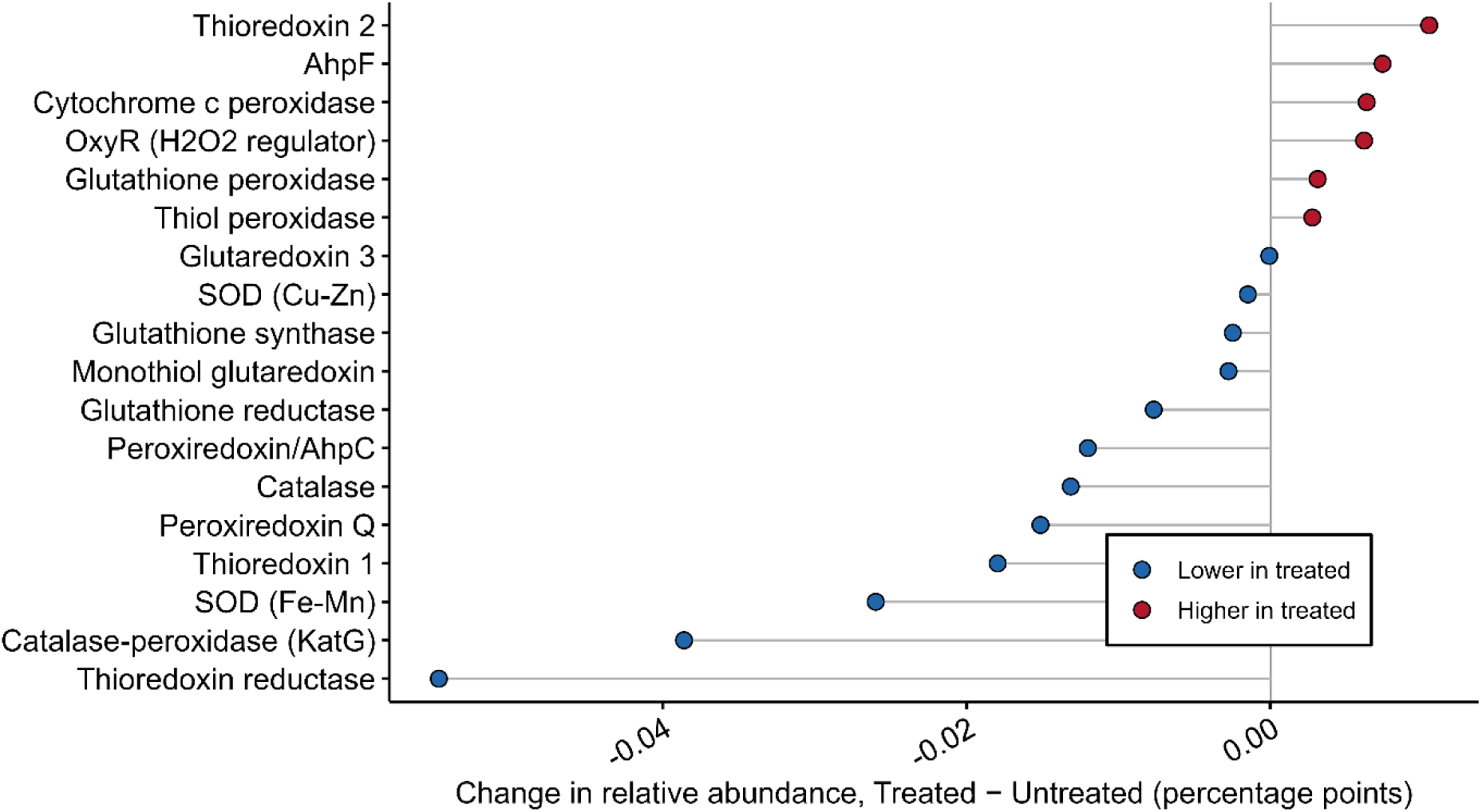
Genomic potential for oxidative-stress-defence gene repertoire shows no coherent enrichment under iDDE treatment. Change in relative abundances (treated-untreated, percentage points) for 18 oxidative-stress-defence orthologs (KEGG level 4), ordered by magnitude of change. Values are means of the three sequential grab samples per conditions; no inferential statistics were applied; points right of zero (red) are higher in treated effluent, left of zero (blue) lower. Most orthologs – including the largest contributors, thioredoxin reductase, catalase-peroxidase (KatG), and Fe-Mn superoxide dismutase – trend lower under treatment, and the panel as a whole is not enriched. Because these profiles derive from DNA, they reflect gene presence in the surviving genomes rather than expression, and are compositionally coupled to the taxonomic shift rather than independent of it. KEGG-assignable read fractions were also markedly lower in treated samples (mean 1.9% vs 4.9%), further limiting DNA-level functional resolution.

## 4. Discussion

By integrating hydrochemistry, stable-isotope analyses and shotgun metagenomics, this pilot study aimed to characterise the short-term effects of BDD treatment on real WWTP effluent beyond conventional reactor-performance metrics. The combined results indicate that treatment affected the bulk matrix only marginally while generating an isotopically distinct oxygen pool consistent with water oxidation and reproducibly restructuring the recovered community DNA, though without a resolvable change in functional gene potential.

### 4.1. Effluent chemistry remains mostly unchanged following treatment

The limited changes observed across the measured physicochemical and geochemical parameters indicate that a 10-min iDDE treatment induced only minor changes to the bulk effluent matrix. Electrical conductivity, alkalinity, the major dissolved ions, as well as the bulk carbon pools remained largely unaffected, suggesting that the treatment did not substantially alter the overall ionic composition or carbon pools of the effluent over the investigated time period. This limited response is notable given the electrochemical conditions applied (7 V, 1.02 A) and suggests that the predominant effects were not translated to bulk water chemistry.

The pronounced decrease in Eh represents the main exception to this overall stability and indicates, if considered in isolation, a shift towards more reducing conditions (e.g., Sigg 2000). However, in an electrode environment, bulk Eh measurements do not necessarily reflect the highly localized redox conditions established at the electrode’s surface during electrolysis (Read and Macpherson 2016; Chen et al. 2024; Huang and Mei 2026). In the present undivided, interdigitated electrode configuration, anodic and cathodic reactions occur within the same solution volume, hence, the measured Eh after treatment actually reflects the products of both half-reactions (Zulla et al. 2025). Moreover, the measurements were executed after the applied current was terminated and therefore represent the post-treatment bulk rather than the transient electrochemical environment at the electrode surfaces. Cathodically generated H2 may further influence the response of the Pt redox electrode (i.e, sensor) and contribute to the observed decrease in bulk Eh (Prass et al. 2021; Qu and Liang 2022; Williams and Lopes 2025). While the measured Eh remains a valuable indicator of the system’s overall post-treatment state, it should not be interpreted as a direct or exclusive measure of the oxidative conditions generated at the anode.

This bulk Eh shift is, to a small extent, reflected in the small changes in dissolved metals and phosphate (Fig. 3), which provide evidence that the treatment actually exerted some influence redox-sensitive components of the effluent. In the untreated effluent, which was oxygenated and near-neutral pH (∼6.7), Fe and Mn would be expected to occur mainly in their oxidised forms, or either be chelated and/or in association with particulate or mineral phases, rather than as freely dissolved reduced species (Davison 1993; Cornell and Schwertmann 2003; Wandersman and Delepelaire 2004). The pronounced decrease in bulk Eh following iDDE treatment therefore raises the possibility of transient reductive processes during electrolysis, including reduction or dissolution of Fe-and Mn-bearing phases (Cornell and Schwertmann 2003; Catrouillet et al. 2020; Makhanbetov et al. 2026). Subsequent mixing with the oxygen-saturated bulk solution most likely favoured reoxidation and reprecipitation, thus resulting in the net changes observed in the dissolved Fe and Mn pools. However, because only dissolved concentrations were determined before and after treatment, and Fe and Mn oxidation state and solid phases were not investigated, this interpretation remains tentative. Nevertheless, the concomitant decrease in phosphate may likewise reflect interactions with Fe-and Mn-bearing phases (Weng et al. 2012; Kraal et al. 2022; Anschutz et al. 2024), although direct proof for sorption or co-precipitation is not available. Thus, rather than indicating straightforward quantitative removal, the Fe and Mn responses are best interpreted as evidence for redox-sensitive cycling during and following iDDE treatment.

A similar low responsivity was observable for the carbon data, thus further emphasizing that the iDDE treatment did not substantially alter bulk effluent chemistry. Changes in DIC, DOC, and POC concentrations were small (Fig. 4), and the corresponding δ^13^C values remained within a narrow range. The slight decrease in DIC, together with the small increase in pH, may reflect changes in the carbonate system during electrolysis. CO2 loss to the gas phase is one plausible contribution, particularly under the vigorous mixing and gas evolution associated with the treatment (Noyes et al. 1996). Furthermore, BDD electrodes have been demonstrated to support electrochemical CO2 reduction (Tariq et al. 2024) and, in turn, harbouring the potential to act as sink for DIC. Yet, based on the present data, these processes cannot be distinguished and the small magnitude for both DIC decrease and pH increase, as well as the nearly unchanged alkalinity, do not support substantial depletion of the bulk DIC pool. The near-absence of change in DOC concentration and δ^13^C-DOC likewise suggest that the short treatment period did not induce the commonly observed extensive transformation or mineralisation of the bulk DOC pool (Panizza et al. 2008; Garcia-Segura et al. 2015). Thus, although carbon-transforming reactions may occur locally at the electrode surfaces, their net effect on the bulk carbon pools over the 10-min treatment period was apparently limited.

The largely unchanged bulk chemistry presented so far, however, does not imply an absence of electrochemical transformation products. The methods used here captured changes in major dissolved constituents and carbon pools, but did not include several potentially relevant secondary products of BDD electrochemistry. In particular, chloride-containing effluents may undergo formation of chlorinated oxyanions, including chlorate and perchlorate, at the BDD anodes (Bergmann and Rollin 2007; Azizi et al. 2011; Lee et al. 2026). Neither chlorate nor perchlorate was quantified in the present study, yet their production is to be expected (Bergmann et al. 2015). The absence of a measurable change in chloride therefore provides only limited evidence against their formation, as conversion of a small fraction of the comparatively large chloride pool could generate environmentally relevant concentrations without producing a detectable change in bulk chloride. Quantification of chlorate and perchlorate is consequently required to fully assess the environmental implications of iDDE treatment and should be included in future evaluations of the system, particularly for discharge-relevant applications.

### 4.2. Depleted δ^18^O-DO points toward electrochemical oxygen generation

In contrast to the, if at all, small changes observed in major ions and carbon pools, δ^18^O-DO decreased substantially following iDDE treatment, providing the strongest evidence in the present dataset for oxygen generation during treatment. DO concentrations determined from headspace equilibration for δ^18^O-DO analysis and measured *in situ* using the WTW multiparameter sensors differed systematically in absolute values but exhibited the same direction of change (Fig. 5). The equilibration-derived concentrations are therefore used here for consistency with the isotope measurements.

The δ^18^O-DO value decreased by 31.3 ±0.28‰, from ∼24.7‰ in the untreated effluent to ∼-6.6‰ after treatment (Fig. 5). The untreated value is consistent with atmospheric equilibrium (+24.6 ±0.4‰; Dordoni et al. 2022), whereas the pronounced shift towards lower values indicates the contribution of an isotopically distinct oxygen source during treatment. Simple equilibration of the initial O2 pool with atmospheric O2 cannot readily account for this shift, particularly given the concurrent, albeit small, increase in DO concentration. Instead, the direction of the isotope shift is consistent with the generation of O2 from water at the anode, as O2 produced via the oxygen evolution reaction (OER) can retain an isotopic composition close to that of the substrate water (e.g., Haschke et al. 2018). The pronounced δ^18^O shift relative to the small net change in DO concentration further indicates substantial turnover of the dissolved O2 pool rather than simple accumulation of newly generated oxygen (Venkiteswaran et al. 2007; Tobias et al. 2007; Kim and Kang 2023).

This contrast between the pronounced shift in δ^18^O-DO values and the relative small net increase in DO seemingly supports that oxygen production was accompanied by concurrent removal from the dissolved pool. The observed increase of 0.16 ±0.03 mmol/L represents only a fraction of the theoretical O2 production based on the applied charge (1.02 A x 600 s), which was ∼0.79 mmol/L assuming four electrons per O2 molecule and 100% current efficiency (see ESM E). This difference most likely reflects competing electrochemical reactions, including the formation of other anodic oxidation products (Bergmann and Rollin 2007; Lee et al. 2026), as well as loss of electrogenerated O2 through gas exchange during vigorous mixing and gas evolution. The small net increase in DO should therefore not be interpreted as an estimate of the amount of O2 generated, but rather as the residual change resulting from simultaneous production and removal. A similar pattern observed in a time-resolved test experiment conducted at lower applied voltage, with δ^18^O-DO progressively approaching the isotopic composition of the source water while DO concentrations remained comparatively stable (SI Fig 1). This progressive adjustment towards the source-water value mirrors the behaviour reported for electrocatalytic oxygen evolution, where isotopic discrimination diminishes as the applied overpotential increases and the reaction becomes less selective, so that the evolved O2 approaches the composition of the substrate water (Haschke et al. 2018). Although this experiment is not used quantitatively here, it additionally supports the hypothesis of substantial turnover of the dissolved O2 pool during treatment. This interpretation is further corroborated by the unchanged isotopic composition of the water itself (Fig. 5). Neither δ^2^H-H2O nor δ^18^O-H2O differed notably between treated and untreated effluent (Δδ^2^H-H2O:-0.1 ±0.3 ‰; Δδ^18^O-H2O: +0.05 ±0.07‰). However, the amount of water involved in O2 generation is negligible relative to the bulk water reservoir and would therefore not be expected to produce measurable change in the water-isotope composition (Guy et al. 1993; Haschke et al. 2018). Consequently, the unchanged water isotopes neither fully support nor contradict a water-derived origin of the generated O2.

Taken together, the pronounced decrease in δ^18^O-DO, its consistency with a water-derived oxygen source, and the small net change in DO indicate substantial oxygen turnover during iDDE treatment. These observations are compatible with anodic water oxidation at the BDD surface and provide evidence for electrochemical O2 generation. However, the δ^18^O-DO signal specifically traces molecular oxygen and does not directly demonstrate the formation of reactive oxygen species (ROS). Although BDD anodes can promote the formation of these highly reactive oxidizing species during water oxidation (De Paiva Barreto et al. 2015; Ganiyu and Martínez-Huitle 2019; Counihan et al. 2019), the present results should be interpreted as evidence for electrochemical oxygen generation and associated oxidative chemistry rather than as direct evidence for ROS formation.

### 4.3. BDD-based electrochemical treatment alters the composition of recovered microbial DNA

Beyond the comparatively limited geochemical response, iDDE treatment was associated with a consistent restructuring of the recovered microbial DNA pool. Treated and untreated samples separated according to treatment at the family level, and the direction of the major taxonomic responses was generally consistent across the three sequential grab samples, although response magnitudes varied between samples (Fig. 6). The largest increase was observed for *Comamonadaceae*, which increased from 12.3% to 25.1% of the recovered sequence reads, followed by *Zoogloeaceae* (3.3% to 8.4%). In contrast, *Hyphomicrobiaceae*, *Methylophilaceae*, *Planctomycetaceae* and *Microbacteriaceae* decreased following treatment. The same general pattern was observed at the genus level, where the increase in *Comamonadaceae* was distributed across several genera, including *Acidovorax* (2.4% to 5.6%), *Malikia* (0.9% to 3.7%), *Variovorax* (0.6% to 2.2%), and *Rhodoferax* (0.9% to 1.9%), rather than being attributable to a single taxon (Fig. 7). Such coordinated shifts across multiple taxa suggest that the iDDE treatment affected the composition of the microbial material recovered after treatment rather than producing a random redistribution of individual taxa.

The observed restructuring is compatible with differential susceptibility of microbial taxa to the conditions generated during BDD-based electrochemical treatment. EAOPs can affect microorganisms through multiple mechanisms, including oxidative damage to cell envelopes, proteins and nucleic acids, with the resulting response depending on the oxidant, exposure conditions and organism-specific properties (Moreno-Andrés et al. 2018; Martínez-Huitle and Brillas 2021; Seixas et al. 2022). Consequently, taxa differing in cell-envelope properties, aggregation behaviour, extracellular polymeric substances (EPS), or intrinsic oxidative-stress defence may not be affected equally (Bruguera-Casamada et al. 2016; Ma et al. 2024; Wang et al. 2025). For instance, Schorr et al. (2019) reported strong reductions in culturable bacteria following BDD treatment of sewage-treatment-plant effluent, with the recoverable culturable community becoming dominated by Gram-positive spore-forming bacteria, whereas Gram-negative bacteria were no longer recovered via cultivation. Likewise, experiments using a laser-structured double-diamond electrode (DDE) demonstrated charge-and time-dependent microbial inactivation, while specific organisms such as *Bacillus subtilis*, *Staphylococcus haemolyticus* and *Streptococcus mitis*, remained recoverable after 5 min of treatment at 6 V (Koch et al. 2020). Furthermore, studies investigating WWTP microbial taxa could also frequently identify members of *Comamonadaceae* and *Zoogloeaceae* in wastewater microbial communities, linking their generally high abundances to their metabolic versatility and, in some cases, tendency to associate with biofilms or flocs (Rosselló et al. 1995; Huang et al. 2022; Wang et al. 2022). Comparable shifts towards β-proteobacterial taxa and *Zoogloea* have also been reported in electrochemical wastewater treatment systems (Tang et al. 2016; Das et al. 2026). Additionally, Di Cesare et al. (2026) reported treatment-associated shifts in wastewater bacterial communities following tertiary treatment, providing further support that microbial response to wastewater treatment does not need to be limited to uniform reductions in abundances. Together, these studies support the interpretation that BDD-based electrochemical treatment can impose strong but taxon-dependent microbial shifts, yet, these observations do not establish that the taxa increasing in relative sequence abundance were intrinsically more resistant to iDDE treatment. Rather, they indicate that the diverse components of the microbial community simply responded differently to the treatment.

This distinction is crucial as the metagenomic profiles describe the composition of DNA recovered after treatment rather than the abundance of viability of intact cells. DNA was extracted from material retained on 0.2 µm filters and may therefore include DNA associated with intact cells as well as DNA originating from damaged or lysed cells. Recent work on activated-sludge treatment has demonstrated that intracellular and extracellular DNA response to wastewater treatment varies, highlighting that treatment-associated changes in DNA profiles do not necessarily correspond directly to changes in viable microbial populations (Sabatino et al. 2026). This represents an important general limitation when interpreting DNA-based community profiles following antimicrobial or oxidative treatment. In the present study, however, the available sequence quality-control data provide no indication of a systematic deterioration of sequencing quality following iDDE treatment. All samples passed quality control, with Q30 values between 93.13 and 94.31% and comparable GC contents between 54.7 and 57.1%, while the number of input reads was also sufficient across all samples (SI Table 2). Thus, although treatment-induced DNA damage cannot be excluded at the molecular level, the observed taxonomic restructuring is unlikely to be explained simply by a general loss of sequencing quality or a pronounced treatment-specific sequencing artefact.

The observed changes therefore most plausibly reflect differences in the microbial material represented in the recovered DNA pool, potentially resulting from differential cell disruption, DNA-persistence, relative abundance or recovery. Importantly, the short treatment period (10 min) was too brief to allow the observed increase in relative sequence abundance to result from microbial growth. This is further corroborated by the time-resolved test experiment (SI Fig. 2), which suggests that electrochemical treatment can cause microbial inactivation under similar conditions (30 min, lower voltage), but it does not resolve the fate of the individual taxa during the 10-min treatment. Consequently, the metagenomic data should not be interpreted as evidence for selective survival. Instead, they indicate that iDDE treatment altered the composition of the microbial material recovered from the effluent, which may thus provide a sensitive indication of treatment effects that is not captured by bulk-water chemistry alone.

An additional consideration is the difference in taxonomic assignment rates between treatments. The proportion of reads assigned to the family and genus levels increased from 59.5% to 67.2% and from 45.1% to 54.5%, respectively, following iDDE treatment. This may partly reflect a shift towards taxa that are better represented in the reference databases and therefore contribute disproportionately to the classified fraction. Such database-dependent effects provide an additional reason to interpret changes in relative sequence abundance cautiously. Nevertheless, the consistency of the major taxonomic responses across the sequential samples and their occurrence at both family and genus level indicate that the microbial component of the effluent responded measurably to iDDE treatment.

### 4.4. Functional gene potential does not mirror the observed taxonomic restructuring

In contrast to the pronounced taxonomic restructuring, the relative abundance of genes associated with oxidative-stress defence exhibited no coherent directional response to iDDE treatment. Of the 18 KEGG orthologs examined, six increased and 12 decreased, which all changes remaining within a narrow range of-0.055 to +0.005 percentage points (Fig. 9). The features showing slightly higher relative abundance included thioredoxin 2, alkyl hyperperoxide reductase subunit F (AhpF), cytochrome *c* peroxidase, OxyR, gluthatione peroxidase and thiol peroxidase, whereas thioredoxin reductase, catalase-peroxidase (KatG), Fe-Mn superoxide dismutase, thioredoxin 1, peroxiredoxin Q and catalase generally decreased.

At first sight, this contrasts with studies reporting pronounced expression of catalases, superoxide dismutases, peroxidases and thioredoxin systems following oxidative treatment (Chen et al. 2019; Sun et al. 2025). However, these studies investigated transcriptional responses, whereas the present analysis characterises functional potential encoded in recovered genomic DNA. A transient transcriptional response during the 10-min treatment would therefore not necessarily result in a measurable change in the relative abundance of the corresponding genes. Conversely, the absence of a metagenomic response does not demonstrate that microorganisms were not exposed to oxidative stress. Simply put, the two types of measurements address different levels of biological response.

The lack of a coherent functional response may additionally reflect the broad distribution of oxidative-stress-defence functions across wastewater-associated bacteria. Catalases, superoxide dismutases, peroxidases and thioredoxin systems occur across diverse bacterial lineages, including taxa commonly associated with activated sludge and wastewater effluent (Wu et al. 2019; Bains et al. 2019; Vestergaard et al. 2024). Taxonomic restructuring may therefore occur without a corresponding change in the overall representation of these functions if different members of the community encode overlapping stress-defence capacities. Such functional redundancy has been proposed as mechanism contributing to the maintenance of community-functional potential despite taxonomic turnover (Louca et al. 2018). In the present study, this interpretation remains tentative because the metagenomic data do not resolve whether individual functional features were retained within viable cells or simply remained represented within the recovered DNA pool.

The short exposure period provides an additional possible explanation for the limited functional response. The 10-min treatment may be sufficient to affect microbial cell integrity and alter the composition of the recoverable DNA pool without producing measurable changes in genomic functional representation. As immediate physiological responses to oxidative stress and subsequent changes in community-level functional potential occur on different temporal scales, the transcriptional responses previously might therefore not be directly translated into altered metagenomic gene abundances of the short-term exposure (Chen et al. 2019; Sun et al. 2025). Moreover, the heterogeneous composition of real WWTP effluent, including organic matter and suspended material (Rajab et al. 2015; Williams et al. 2019), may create microscale differences in oxidant exposure that further limit a uniform community-wide response.

Nevertheless, the functional data should be interpreted together with the sequencing QC rather than attributed to a general deterioration of DNA quality. The proportion of reads assigned to KEGG orthologs was lower in treated than in untreated samples (1.9% versus 4.9%), which could in principle result from treatment-associated DNA damage or differences in DNA recovery. EAOPs are known to cause oxidative damage to nucleic acids, which can involve strand beaks and base modifications, thus increasing the risk of compromising the recovery and subsequent sequencing of the affected DNA (Costello et al. 2013; Child et al. 2023; Wang et al. 2024). However, considering the consistently high Q30 values, absence of QC failures and comparable GC contents across all samples, a broad treatment-related degradation of sequencing quality seems unlikely. The lower KEGG-assigned fraction may instead reflect differences in taxonomic composition, database representation or recoverability of particular DNA fragments. Consequently, DNA damage remains a mechanistically plausible effect of oxidative treatment, but the present QC data do not support it as the primary explanation for the observed functional profiles.

Taken together, the metagenomic results indicate that short-term iDDE treatment altered the composition of microbial DNA recovered from real WWTP effluent without producing a corresponding change in the overall representation of the oxidative-stress-defence genes examined here. This apparent decoupling between taxonomic composition and functional potential does not indicate the absence of microbial stress, nor does it demonstrate selective survival. Rather, it demonstrates that the microbial response to iDDE treatment can be detected at the level of community composition even when bulk chemistry and the examined genomic functional potential remain comparatively stable. This suggests that microbial community profiling may provide an additional dimension for evaluating EAOP treatment effect, while also highlighting the need for complementary measurements capable of distinguishing inactivation, physiological stress and changes in DNA persistence.

## 5. Conclusions and Outlook

This pilot study demonstrates that short-term BDD-based treatment of real WWTP effluent can induce measurable electrochemical and microbial response despite comparatively small changes in bulk water chemistry. The strongly depleted δ^18^O-DO values, together with the small net increase in dissolved oxygen, are consistent with substantial turnover of the dissolved oxygen pool and provide evidence compatible with electrochemical oxygen generation at the BDD electrode. Shotgun metagenomics further revealed pronounced taxonomic restructuring of the recovered microbial DNA pool. Comparison with previous culture-based BDD treatment of wastewater and DDE-mediated microbial inactivation indicates that heterogenous microbial responses to diamond-electrode electrochemistry are indeed plausible, however, the present DNA-based data cannot distinguish microbial inactivation from DNA persistence, differential cell disruption or differential recovery. Likewise, the absence of a coherent response in the oxidative-stress-defence gene repertoire does not exclude physiological stress, but highlights the distinction between genomic functional potential and an immediate microbial stress response. Yet, sequencing QC did not indicate a general treatment-related loss of data quality, thus suggesting that the observed community restructuring was not simply attributable to sequencing artefacts.

Overall, these findings indicate that the consequences of BDD-based electrochemical treatment extent beyond bulk-water chemistry and conventional measures of microbial inactivation. At the same time, the results should not be interpreted as evidence for selective survival or taxon-specific resistance. Resolving the mechanisms underlying the observed microbial community restructuring will require complementary measurements of absolute microbial abundance and viability, intracellular versus extracellular DNA, and microbial activity. Such measurements would allow future studies to determine whether the compositional shifts observed here reflect differential inactivation, persistence of extracellular DNA, or genuine differences in microbial resistance to BDD-generated oxidative conditions.

## 6. Data availability

The dataset for hydrochemical and isotope analyses is provided as ESM, whereas the metagenomic sequences are to be deposited at NCBI.

## 7. Author contributions

ANV conceptualized and designed the study, acquired funding, performed sampling, laboratory work and data analyses, prepared the figures, and wrote the manuscript.

## 9. Acknowledgements

The authors sincerely thank the Stadtentwässerung und Umweltanalytik Nürnberg (SUN) for facilitating access to the WWTP I in Nürnberg-Muggenhof. Special thanks are due to Karin Leuthold-Schmitt, Claudia Reitz, Andreas Seidel and Hartmut Klügel for their essential support, and to the PhD students Christina M. Schubert and Aixala Gaillard for their dedication during the sampling campaign. We also gratefully acknowledge Prof. Dr. Kathrin Castiglione and Matthias Tobler for providing the Qubit system and assisting with its operation. We also thank Prof. Dr. Johannes Barth for providing the laboratory infrastructure, consumables, and technical facilities that supported this independent research. In this regard, we also like to thank Irene Wein and Christian Hanke for their contributions. Additionally, we thank Dr. Marcus Speck for the dry ice rescue. Finally, we would like to thank Maxima Praxl for her valuable work during her Bachelor’s thesis, which provided data used in the supplementary materials.

During the preparation of this manuscript, the author used Claude (Opus 4.8; Anthropic) for refinement and annotation of R code used for data visualisation. The author reviewed and verified all outputs and takes full responsibility for the content of the manuscript.

## 10. Funding resources

This study was funded by the “Dr. Hertha und Helmut Schmauser-Stiftung” of the Friedrich-Alexander-University Erlangen-Nürnberg.

