## Supplementary Information for "Short-term BDD treatment of wastewater effluent reveals dissolved oxygen turnover and microbial community restructuring"

ANV: 0000-0002-8447-5825

MZ:

SR: 0000-0002-5728-0635

AB: 0000-0003-1896-4521

### Supplementary Figures

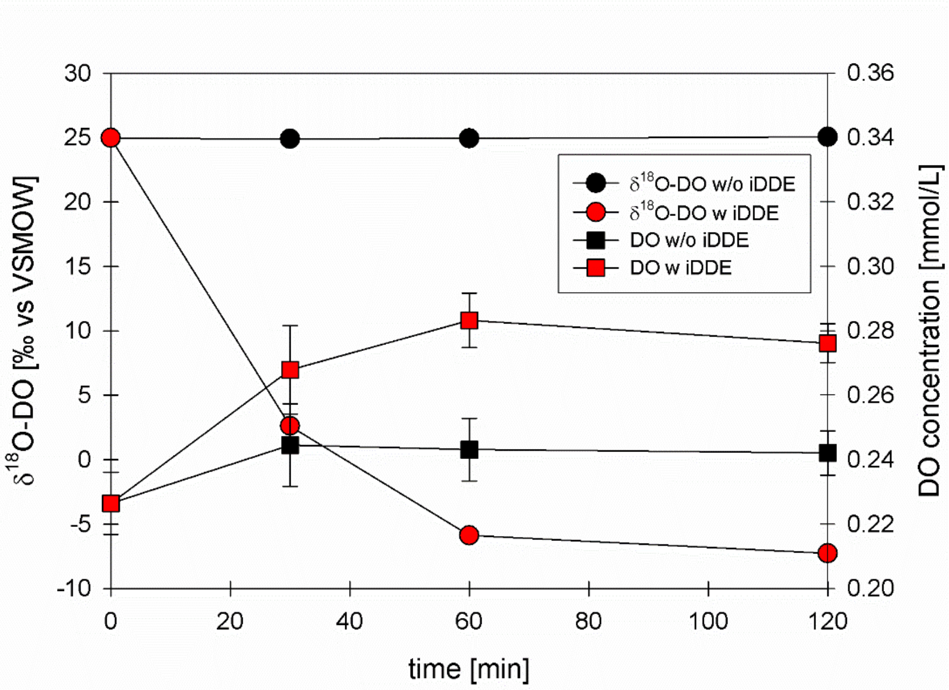


*Figure S 1:* Time-resolved response of dissolved oxygen concentration and $\delta^{18}$O-DO of effluent treated with (red) and without (black) the interdigitated double diamond electrode (iDDE). Unlike the main experiment, this test used effluent water from the Erlanger EBE WWTP and followed $\delta^{18}$O-DO (dots) and DO concentrations (squares) for 120 min at 6 V and 0.56 A. Note different scaling of the y-axis. Error bars represent ±SD ($n$ = 3).

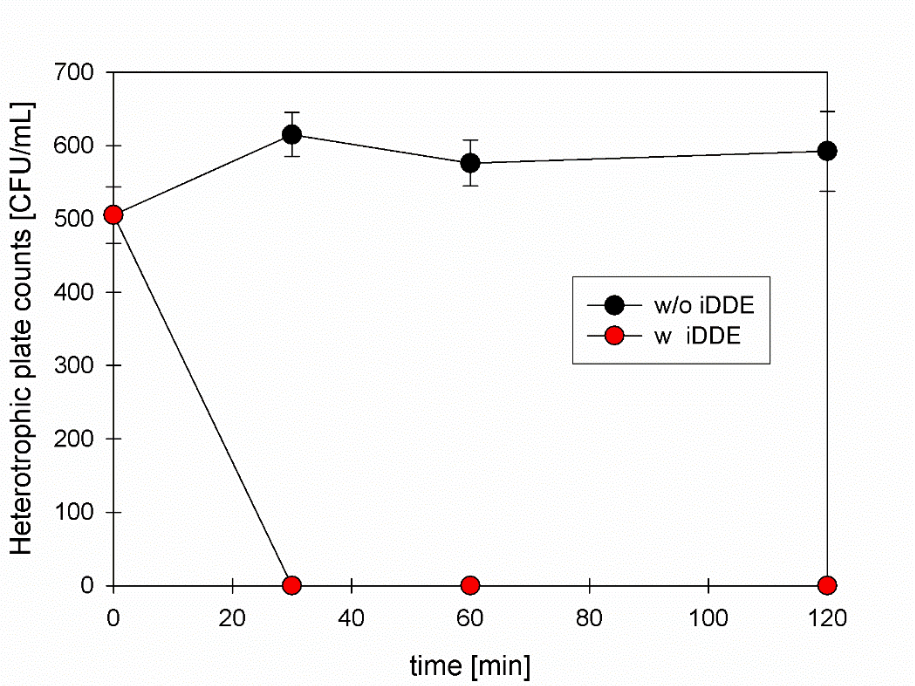


*Figure S 2:* Time-resolved cultivable heterotrophic bacteria numbers in effluent treated with (red) and without (black) the interdigitated double diamond electrode (iDDE). Heterotrophic plate counts (CFU/mL) in effluent from the Erlanger EBE WWTP over a treatment duration of 120 min (6 V, 0.56 A), determined using the IDEXX Easy Disc DEV Test Kit incubated at 20°C for 44 hrs. Error bars represent ±SD ($n$ = 3).

### Supplementary Tables

*Table 1:* DNA concentration of effluent samples determined by fluorometric quantification compared to values reported by the sequencing provider (CeGaT, Tübingen, DE). Genomic DNA concentrations (ng/µL) recovered from untreated (WWTP_2,6,10) and iDDE-treated (WWTP_4,8,12) effluent samples, quantified in-house using the Qubit dsDNA BR Assay Kit (Invitrogen) and as reported by CeGaT during library quality control prior to sequencing. Values represent the mean of *n* = 3 measurements per sample)

| Sample ID | Sample volume µL | Qubit BR kit [ng/µL] | CeGaT [ng/µL] |
|---|---|---|---|
| WWTP_2 | 1 | 4.76 | 7.6 |
| WWTP_6 | 1 | 6.52 | 1.4 |
| WWTP_10 | 1 | 0.88 | 9.6 |
| WWTP_4 | 1 | 11.8 | 16.1 |
| WWTP_8 | 1 | 3.04 | 4.5 |
| WWTP_12 | 1 | 37.7 | 51.9 |

*Table 2:* Sequencing statistics and quality-control metrics for shotgun metagenomic libraries. Raw sequencing statistics and quality-control metrics for untreated (WWTP_2,6,10) and iDDE-treated (WWTP_4,8,12) effluent samples, as reported by CeGaT (Tübingen, DE) on the Illumina DRAGEN Bio-IT Platform (v4.4.4): Number of input reads, percentage of reads failing quality control (QC fail %), unaligned reads (unmapped %), duplicated reads (Dup %), properly paired reads (Prop pair %), based with Phred quality score ≥30 (Q30 %) and GC content (%GC).

| Sample ID | M Input reads | QC-fail % | Unmap % | Dup % | Prop pair % | Q30 % | % GC |
|---|---|---|---|---|---|---|---|
| WWTP_2 | 48.5 M | 0 | 100 | 0 | 0 | 93.33 | 56.57 |
| WWTP_6 | 89.6 M | 0 | 100 | 0 | 0 | 94.31 | 56.92 |
| WWTP_10 | 53.9 M | 0 | 100 | 0 | 0 | 93.6 | 54.67 |
| WWTP_4 | 48.3 M | 0 | 100 | 0 | 0 | 93.13 | 57.08 |
| WWTP_8 | 59.5 M | 0 | 100 | 0 | 0 | 94.11 | 56.83 |
| WWTP_12 | 52.3 M | 0 | 100 | 0 | 0 | 93.71 | 54.91 |
